# Fine-scale spatial patterns of wildlife disease are common and understudied

**DOI:** 10.1101/2020.09.01.277442

**Authors:** Gregory F Albery, Amy R Sweeny, Daniel J Becker, Shweta Bansal

## Abstract

1. All parasites are heterogeneous in space, yet little is known about the prevalence and scale of this spatial variation, particularly in wild animal systems. To address this question, we sought to identify and examine spatial dependence of wildlife disease across a wide range of systems.
2. Conducting a broad literature search, we collated 31 such datasets featuring 89 replicates and 71 unique host-parasite combinations, only 51% of which had previously been used to test spatial hypotheses. We analysed these datasets for spatial dependence within a standardised modelling framework using Bayesian linear models, and we then meta-analysed the results to identify generalised determinants of the scale and magnitude of spatial autocorrelation.
3. We detected spatial autocorrelation in 48/89 model replicates (54%) across 21/31 datasets (68%), spread across parasites of all groups. Even some very small study areas (under 0.01km^2^) exhibited substantial spatial variation.
4. Despite the common manifestation of spatial variation, our meta-analysis was unable to identify host-, parasite-, or sampling-level determinants of this heterogeneity across systems. Parasites of all transmission modes had easily detectable spatial patterns, implying that structured contact networks and susceptibility effects are potentially as important in spatially structuring disease as are environmental drivers of transmission efficiency.
5. Our findings demonstrate that fine-scale spatial patterns of infection manifest frequently and across a range of wild animal systems, and many studies are able to investigate them whether or not the original aim of the study was to examine spatially varying processes. Given the widespread nature of these findings, studies should more frequently record and analyse spatial data, facilitating development and testing of spatial hypotheses in disease ecology. Ultimately, this may pave the way for an *a priori* predictive framework for spatial variation in novel host-parasite systems.

## Introduction

The maintenance and spread of parasites are inherently spatially structured (Cross, Lloyd-Smith, Johnson, & Getz, 2005; Kirby, Delmelle, & Eberth, 2017; Pullan, Sturrock, Soares Magalhaes, Clements, & Brooker, 2012), which holds important ramifications for epidemiological dynamics and disease control efforts (Becker et al., 2020; Cross et al., 2005; Plowright, Becker, McCallum, & Manlove, 2019). Spatial structure can arise through a wide variety of processes: for example, many parasites are transmitted from one host individual to another via direct contact, which requires a degree of spatiotemporal coincidence between individuals (Manlove et al., 2018), so that infections are spatiotemporally staggered in waves of transmission across the population. Other parasites transmit through persistent environmental stages or arthropod vectors whose viability depends on spatially varying abiotic conditions, creating spatial patterns of exposure and therefore of infection (Altizer et al., 2006; Jamison, Tuttle, Jensen, Bierly, & Gonser, 2015; Patz, Graczyk, Geller, & Vittor, 2000). Finally, host immunity and susceptibility can be influenced by environmentally varying factors like resource availability and climatic conditions, with knock-on impacts on parasite burden and transmission (Becker et al., 2020, 2018). These diverse processes should produce spatial patterns of infection across a wide range of wildlife systems, yet many wildlife disease studies examine coarse spatial scales or assume that spatial patterns will be negligible compared to other hypothesised drivers. As such, it is unclear how often infection is spatially structured in these systems, at what range this variation can manifest, and how host and parasite traits might alter its manifestation.

For logistical reasons, many studies of spatial drivers of infectious disease focus on discrete between-population differences across large distances, often using a limited number of discrete sampling locations rather than distributing their sampling locations continuously in space (Plowright et al., 2019). Nevertheless, recent work suggests that spatial patterns of infection may manifest at surprisingly fine spatial scales, within kilometres or even metres (Abolins et al., 2018; Albery, Becker, Kenyon, Nussey, & Pemberton, 2019; Brooker et al., 2006; Wood et al., 2007). This observation begs the question: what is the lower bound for the range at which spatial effects can act? Identifying the range of spatial dependence (or autocorrelation, meaning that data points that are closer together in space tend to be more similar) is important for many reasons, including designing sampling regimes (Nusser, Clark, Otis, & Huang, 2008; Plowright et al., 2019; Vidal-Martínez, Pech, Sures, Purucker, & Poulin, 2010), building mechanistic models of parasite evolution over space (Best, Webb, White, & Boots, 2011; Débarre, Hauert, & Doebeli, 2014), examining how disease risk responds to anthropogenic activities like urbanisation (Saito & Sonoda, 2017), and directing public health and conservation schemes (Brooker et al., 2006; Gilbertson et al., 2016).

Identifying the range of spatial dependence can also help examining how parasites spread over landscapes and determining their transmission mechanisms (Reynolds, 1988). For example, spatial dependence across large distances might suggest the influence of major climatic correlates, while spatial dependence between nearby locations suggests a highly localised infection process (Pullan et al., 2012). In human disease systems, such work has shown that neighbouring districts of Thailand have more similar human malaria incidence, suggesting local similarities in abiotic conditions or vector control programs that could limit mosquito survival (Zhou et al., 2005). Similar analyses of wildlife disease could help pinpoint transmission routes and guide disease control effort: for example, if researchers find that a zoonotic disease has a long range of dependence in its wildlife reservoir, this could motivate the use of widely placed sampling locations when trying to identify environmental drivers (Becker, Crowley, Washburne, & Plowright, 2019; Plowright et al., 2019). Lastly, the scale of spatial dependence has implications for more general theoretical understanding of infectious disease dynamics. For example, links between biodiversity and disease dynamics (e.g. “dilution effects”) are dependent on the spatial scale of sampling (Cohen et al., 2016; Rohr et al., 2020), and several rodent systems have identified contrasting spatial trends for zoonotic diseases dependent on sampling scale (Luis, Kuenzi, & Mills, 2018; Morand et al., 2019).

The strength and range of spatial dependence are also likely to depend on the traits of the hosts and parasites involved. For example, parasites that persist for longer in the environment are likely to experience stronger influences of environmental gradients than directly transmitted counterparts (Satterfield, Altizer, Williams, & Hall, 2017). Similarly, highly mobile species such as large carnivores or nomadic bats may more efficiently disseminate parasites through the environment, reducing spatial autocorrelation (Gilbertson et al., 2016; Peel et al., 2013). However, the relative contribution of host and parasite traits to shaping spatial variation in infection remains unknown. The range of spatial dependence is most commonly identified using spatial autocorrelation models (e.g. Albery et al., 2019; Becker, Nachtmann, et al., 2019; Brooker et al., 2006; Gilbertson et al., 2016; Wood et al., 2007) or analyses that quantify the spatial buffer regions in which environmental variables are best-correlated with disease (e.g. Saito & Sonoda, 2017). Unfortunately, these approaches are almost always reactive rather than proactive and they occur on a case-by-case basis rather than being based on general rules or *a priori* understanding. To establish general factors influencing the scale of spatial dependence in wildlife disease, a variety of host-parasite systems must be analysed using comparable techniques and then synthesised. As well as revealing fundamental drivers of spatial heterogeneity, identifying general rules in this way could facilitate the development of predictive models for spatial structuring in host-parasite systems with relatively poorly understood epidemiology (Gilbertson et al., 2016).

Researchers could then predict how within- and between-population processes will differ *a priori*, before using empirical methods such as long-term studies at multiple scales (e.g. Luis et al., 2018; Morand et al., 2019).

Prescriptive rules for examining geographic variation in wildlife disease are rare and hard to generalise. For example, where studies seek to quantify the impact of environmental drivers on parasitism, larger study extents may allow sampling the widest range of different environmental factors and thus increasing spatial variation (Becker et al., 2020; Cohen et al., 2016). Part of this methodological vacuum for generalisable rules emerges from the analytical complexity of identifying them. A recent systematic review of ecoimmunology studies uncovered a surprising lack of spatial methods, with most studies fitting discrete fixed or random effects to control for spatial autocorrelation rather than directly examining continuous patterns in space or using spatially explicit statistics (Becker et al., 2020). Although the statistical competence of ecologists is high and increasing, particularly with regards to areas like movement ecology and network analysis (Albery, Kirkpatrick, Firth, & Bansal, 2021; Dougherty, Seidel, Carlson, Spiegel, & Getz, 2018; Jacoby & Freeman, 2016; Webber & Vander Wal, 2019), no empirical framework exists for establishing the presence or range of spatial variation in wildlife disease. Establishing general factors shaping spatial variation across wildlife disease systems could lay the basis for this framework, improving mechanistic understanding of parasite transmission, spatial sampling designs, and control efforts.

Here, we conducted a synthesis of spatially distributed wildlife disease datasets across a wide range of different host and parasite taxa, geographic contexts, and sampling regimes. We analysed these datasets individually using a standardised modelling procedure, identifying how generalised host-, parasite-, and sampling-level factors affect the prevalence and range of spatial dependence. Specifically, we expected that studies would be most vulnerable to strong spatial effects in larger study areas, with greater sampling efforts, and when parasites exhibit indirect transmission mechanisms with extended environmental stages. We aimed to provide important general estimates for the range of spatial autocorrelation from a wide range of different host-parasite systems, laying the groundwork for *a priori* predictions about host-parasite systems with unknown spatial properties.

## Materials and methods

### Data collection

To obtain a wide set of raw datasets we carried out a literature search, emailed authors to request data, and searched data repositories for publicly available datasets (Supplementary Figure 1). Our literature search used Web of Science to identify potential datasets published between 2009 and 28^th^ August 2019, with the following terms: “(parasit* OR infect* OR disease) AND (wild OR natural) AND (mammal)”. We restricted the search to mammals to increase the generalisability of our findings within this group of animals, and because of their importance for human and livestock health (Han, Kramer, & Drake, 2016).

We first screened a random subset of studies based on their abstracts, excluding studies of captive animals, review papers, and meta-analyses; publications without parasite data; studies without hosts (i.e., only sampling parasites in the environment; and studies of non-mammals. Because our downstream analyses relied upon a standard spatial modelling procedure, we also excluded studies with few samples (N<35), very low prevalence (<10%), or very high prevalence (>90%), owing to likely failure in model convergence.

If a study had openly available datasets we downloaded them, and for those that included binary infection data in map figures, we derived approximate spatial locations and associated infection status (i.e., “heads up digitisation”, HUD). We also searched the Dryad data repository (https://datadryad.org) using the same search terms to find publicly available datasets.

For all other studies, we contacted corresponding authors using a standardised email template in September-December 2019 to request data. We classified the authors’ responses into the following categories (Supplementary Figure 1): System not suitable: the system was poorly suited to our questions (e.g., migratory host population). No parasitology: the system did not include disease measures. No spatial data collected: no sources of spatial data (grid references, GPS locations) were collected and associated with individuals or samples. Privacy concerns: researchers were unable to share the data because they were collected on private land. Data not suitable: once data were inspected, the genre of spatial data was found to be unsuitable (e.g. too few spatial replicates), or it was deemed unlikely that models would run (e.g., points very unevenly distributed, sample sizes too low).

Some of the datasets contained multiple spatial sites that were each defined as a contiguous population. Therefore, within the datasets, each replicate was defined as a unique host-parasite-locality combination examining a contiguous population. We excluded replicates with under 100 samples, to ensure convergence of our standardised spatial models (see below).

Although we principally aimed to quantify fine-scale, within-population spatial effects, we included several studies employing continuous or semi-continuous sampling at national and county levels, to investigate whether the methods we used would operate well at these scales and to establish an upper bound for sampling effects.

### Statistical Analysis

#### Data standardisation

Data were manipulated and analysed using R version 3.6.3 (R Development Core Team, 2011). All code is available at github.com/gfalbery/libra. Our data cleaning procedure aimed to minimise the probability of false positives and to restrict the data pool to a continuous spatial distribution of samples. All spatial coordinates were converted to the scale of kilometres or metres to allow comparison across systems. We removed spatial outliers and parasite count outliers; if parasite counts were very overdispersed and/or highly zero-inflated they were analysed as binomial (0/1) infection data rather than negative binomial. Categories with low replication (generally <10 samples) were removed. We removed specific classes that exhibited very low prevalence: e.g., adult Soay sheep and red deer had a very low prevalence of *Nematodirus* sp., which is primarily a parasite of young ungulates (Hoberg, Kocan, & Rickard, 2001); hence only lambs/calves were analysed. Individual identity was fitted as a random effect if the dataset involved repeat measurements of the same individuals.

#### INLA Models

We based our analysis on a framework previously used in a study of spatial patterns of disease in wild red deer (Albery et al., 2019). Integrated Nested Laplace Approximation (INLA) models were fitted to each spatial dataset using the `inla` package. INLA is a deterministic Bayesian algorithm that allows fitting of a Stochastic Partial Differentiation Equation (SPDE) random effect to quantify and control for patterns of the response variable in space. This relies on detection of spatial autocorrelation, where samples closer in space are more similar than those further apart (Kirby et al., 2017; Tobler, 1970). The model estimates how much variance is accounted for by autocorrelation, and models with and without the SPDE effect can be compared to assess how it affects the fit of the model (Lindgren & Rue, 2015; Zuur, Ieno, & Saveliev, 2017). The model also provides a “range” parameter, which estimates the distance at which samples are autocorrelated. We took this parameter to represent a combination of sampling, transmission, and immune processes determining the scale of spatial variation in the focal population.

We first fitted a “base” model with parasite burden (Gaussian or negative binomial) or presence/absence (binary) as a response variable and with any fixed and random covariates. To simplify our analyses, covariates usually included only temporal variables (month, year, both as categorical variables), age category, and sex. We then fitted a model featuring an SPDE random effect, with a penalised complexity prior (Fuglstad, Simpson, Lindgren, & Rue, 2019). We compared the base model with the SPDE model, identifying whether the latter had a lower Deviance Information Criterion (DIC), indicating improved model fit. We took a change in DIC (ΔDIC) of 2 to distinguish between the two models and calculated the DIC weight for the base and SPDE model, giving a proportion (0-1) that can be conceptualised as “confidence that the spatial model was the best-fitting” (Wagenmakers & Farrell, 2004). We also extracted the INLA range parameters. In total, we fitted INLA models to 89 host-locale-parasite combinations, each of which comprised a different host-locale-parasite combination, generated from 31 different study systems.

#### Meta-analysis of INLA models

To identify factors driving general trends of spatial variation, we conducted a meta-analysis treating each unique parasite-system-site combination as a replicate, including parasite-, host-, and sampling-level traits as fixed effects. We constructed hierarchical models using the ‘metafor’ package. Generally, meta-analyses typically focus on synthesizing effect sizes and their variances across multiple systems (e.g. Sánchez *et al*. 2018). However, as generalised spatial variation does not have a directional effect, we instead analysed measures of model fit, predictive capacity, and the autocorrelation range, which is bounded at 0 and infinity. To give a coarse measure of model predictive capacity that was easily standardised across all models, we calculated the Spearman’s Rank correlation between the observed and predicted values for the model, using only the SPDE effect to predict (henceforth referred to as R). The measures of model fit give an impression of the detectability and importance of spatial patterns, while comparisons of the range estimate across systems will inform whether different host and parasite traits cause spatial patterns to vary more sharply in space. We used the *escalc* function to derive sampling variances for DIC weight and the INLA range (using the point estimate and 95% confidence interval).

Our hierarchical models included each replicate nested within study as a random effect to account for within- and between-study heterogeneity (Konstantopoulos, 2011). We also included a random effect for host family, for which the covariance structure used the phylogenetic correlation matrix (Nakagawa & Santos, 2012); we obtained our phylogeny from the Open Tree of Life with the rotl and ape packages (Michonneau, Brown, & Winter, 2016; Paradis, Claude, & Strimmer, 2004). All models used the `rma.mv` function and weighting by sampling variance. We first assessed heterogeneity in each of our response variables by fitting a random-effects model (REM; intercept only) with restricted maximum likelihood and then used Cochran’s *Q* to test if such heterogeneity was greater than expected by sampling error alone (Borenstein, Hedges, Higgins, & Rothstein, 2009).

We next used mixed-effects models (MEMs) to test how sampling-, host-, and parasite-level factors affected our INLA model outputs. Sampling variables included: Number of samples; Sampling area (total rectangular extent between the furthest points on the X- and Y-coordinates, in km^2^); Sampling method (3 levels: trapping, censusing, and necropsy/convenience sampling); Spatial encoding method (4 levels: GPS; trapping grid; locality; Easting/Northing); Spatial hypothesis testing (binary – i.e., did the study aim to quantify spatial variation in some way?). We interpreted this latter variable as a combination of study design and publication bias, where studies that are intended to pick up spatial variation are both more likely to identify spatial patterns because of their sampling design, and then more likely to be published if they do. Parasite traits included Transmission mode (4 levels: direct; faecal-oral; vector-borne; environmentally transmitted) and Taxon (8 levels: arthropod, nematode, trematode, cestode, protozoan, bacterium, virus, other). Host traits included: Home Range size (in km^2^; log-transformed); Body Mass (in grams; log-transformed); Host order (5 levels: Carnivora, Chiroptera, Ungulates, Glires, Proboscidea). There was only one lagomorph, so rodents and lagomorphs were lumped together into the “glires” clade. The same was true of odd-toed ungulates (Perissodactyla), so they were lumped with Artiodactyla into an “ungulates” clade. For species for which a phenotypic measure (e.g. body mass) was unavailable, we used the value for the closest relative for which the data were available, according to a mammalian supertree (Fritz, Bininda-Emonds, & Purvis, 2009).

To identify important drivers among these many correlated drivers, we conducted a model addition process using maximum likelihood and Akaike Information Criterion corrected for sample size (AICc) to determine model fit. Each of our meta-analytical explanatory variables was added in turn, and the best-fitting variable (i.e., the one that most decreased AICc) was kept for the following round. This process was repeated with the remaining variables, until no variables improved model fit by more than 2 AICc. We report the final model, with the minimal number of variables that improved model fit.

#### Spatiotemporal INLA models

Finally, we constructed spatiotemporal INLA models to assess the consistency of spatial hotspots from year to year, and to investigate evidence of ephemeral waves of transmission across the study systems. Of our 89 replicates, 44 replicates had more than one year of sampling, with more than 100 spatial points per year, facilitating fitting spatiotemporal models. For these replicates, we first reran the original models with the reduced dataset that only included years with more than 100 replicates. We then fitted a spatiotemporal model with a different spatially distributed effect (i.e. “spatial field”) for each year, with no autocorrelation between the fields. Improved model fit for this model would imply that the spatial distribution of the parasite varied notably from year to year. Second, we fitted a similar spatiotemporal model with an “exchangeable” autocorrelation specification between years. This model format allows correlation between spatial fields, but without enforcing a time sequence: that is, all fields were correlated by the same parameter (“Rho”) regardless of how far apart in time they were. The Rho parameter, which is bounded between −1 and 1, was then interpreted to give an impression of the spatiotemporal consistency of the parasite distribution. Parasites with high rho coefficients had very similar hotspots from year to year, while those with low coefficients did not.

## Results

Our literature review returned 3399 studies, and we screened a random selection of 1993 abstracts (over two weeks) to expedite data collection. 1151 of these were unsuitable because they were in the wrong environment, host, or subject area, or had no data. This left 496 studies, for which we assessed data availability and emailed 432 authors to request data if unavailable (Supplementary Figure 1).

Very few studies publicly archived continuous, within-population spatial data. Only 3/496 studies (0.6%) had such data ready to download, and 4 further studies had maps of samples from which we could easily digitise sufficient data (Supplementary Figure 1). We also already owned 3 datasets. When we emailed the corresponding authors of the remaining studies, 92 responded, 22 of which (23.9%) indicated that they had not collected any within-population spatial data as part of their study (Supplementary Figure 1). After navigating a number of other obstacles to data sharing, followed by initial data triage, 26 authors kindly offered to provide us with spatial data, resulting in 36 total viable datasets. Of these 36 datasets, 31 had at least one contiguous spatial population with >100 samples to which we could apply INLA models.

Most authors that responded (and had collected spatial data) were happy to share data with us, and the vast majority of studies for which we did not receive data were due to a lack of response or secondary response (Figure 1). 15 authors responded but declined to share data due to privacy concerns, ongoing data usage, or authorship concerns. Comparing this to the 22 responders that had not collected spatial data implies that the main reason researchers do not share spatial data is that they did not collect it; however, given that >300 researchers did not respond (and they may not have been a random subset of the total), our ability to infer this confidently is diminished. Notably, studies that investigated spatial variation tended to be larger than those that did not (Supplementary Figure 2), implying that larger study areas motivate researchers to more often consider spatial variation in their analyses.

**Figure 1:**
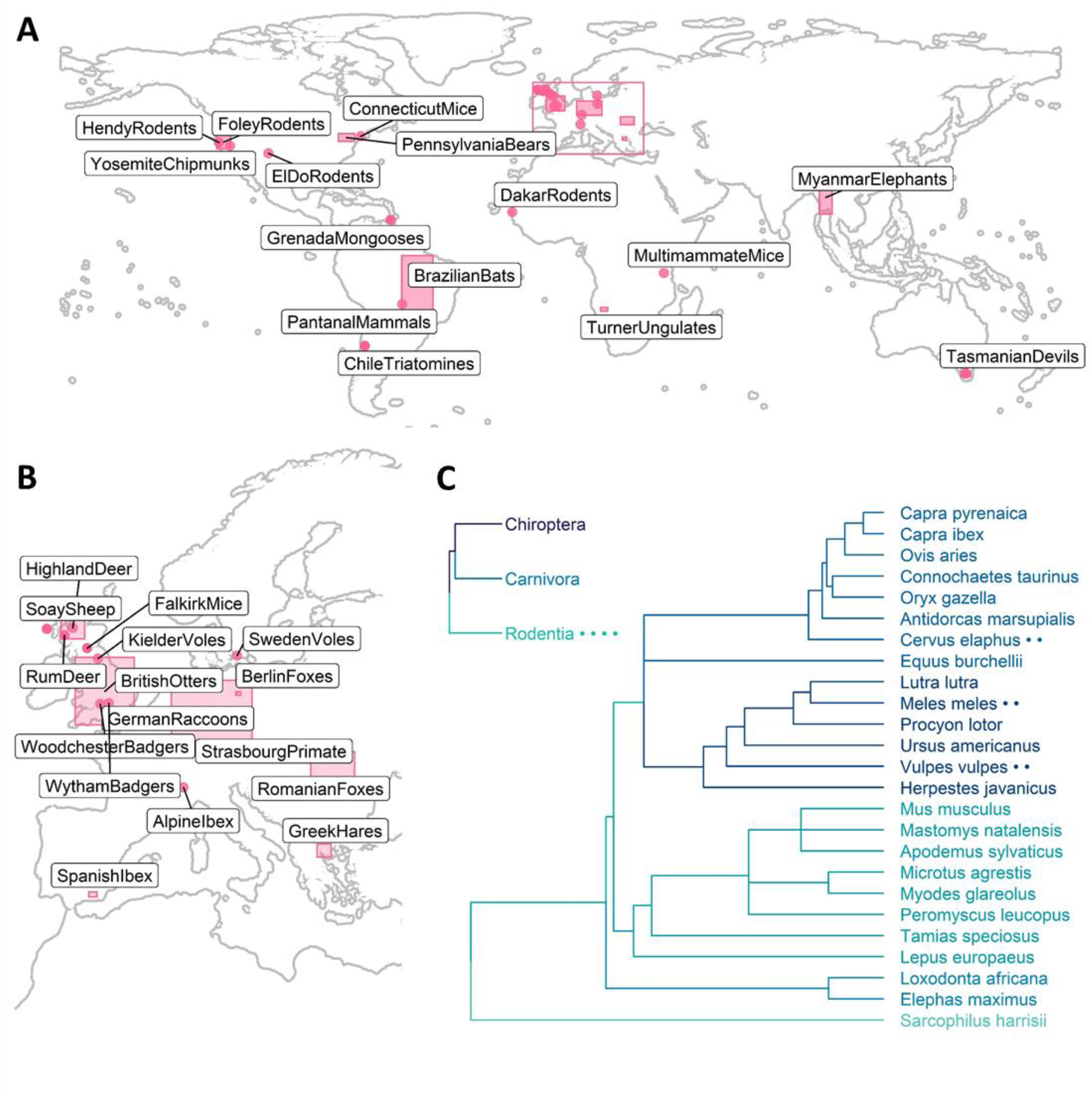
The geographic and taxonomic distribution of the 31 datasets that we included in our final meta-analysis. Our data were evenly spread across the earth (Panel A), although with a notable cluster in Western Europe (see inset map in pink rectangle, Panel B). Sampling areas greater than 5000 km^2^ are displayed as rectangles; smaller sample areas are represented by dots. Study system names correspond to the names in Supplementary Table 1. The datasets also included a wide range of different mammal orders and families (Panel C). The inset phylogeny represents order-level summaries for studies that were not carried out at the species level. Dots next to species’ names in the phylogenies denote that multiple datasets included samples from that species. Different colours correspond to different taxonomic groups used for meta-analysis: ungulates, carnivores, glires, elephants, and carnivorous marsupials.

We concluded data collection with 31 datasets, including 89 spatial replicates and 90 host species (Figure 1). 67 replicates were species-level; the rest were conducted on selections of species in the same order (e.g., rodent trapping, bat sampling, carnivore faecal sampling). The datasets were distributed across five continents (Figure 1), and included 7 different mammalian orders (Figure 1). The studies examined 41 different parasites, across a diverse selection including viruses (N=6), bacteria (N=10), helminths (N=25), arthropods (N=14), and one transmissible cancer (N=8). Infection measures included counts of parasites or immune markers (N=30), binary assessment of infection status using observation or seropositivity (N=52), and one study used parasite-associated mortality as a proxy (Myanmar elephants, *Elephas maximus* (Lynsdale, Mumby, Hayward, Mar, & Lummaa, 2017)). Study systems included, for example: rodent trapping studies examining flea burdens and their associated parasites (e.g. rodents trapped in the Arizona hills (Kosoy et al., 2017) and chipmunks in Yosemite National Park (Hammond et al., 2019)); long-term studies with parasite data collected over the course of several decades (e.g. the Soay sheep of St Kilda (Hayward et al., 2014), the Isle of Rum red deer (Albery et al., 2019), and the badgers of Wytham Wood (Albery et al., 2020)); and studies examining seropositivity of mammals across a geographic range to identify endemic areas (e.g. British otters infected with *Toxoplasma gondii* (Smallbone et al., 2017)). See Supplementary Table 1 for a description of each study system and the associated references and researchers that provided us with the data. The area of the study systems varied widely, from 0.02 to 10^6^ km^2^ (Figure 2A).

**Figure 2:**
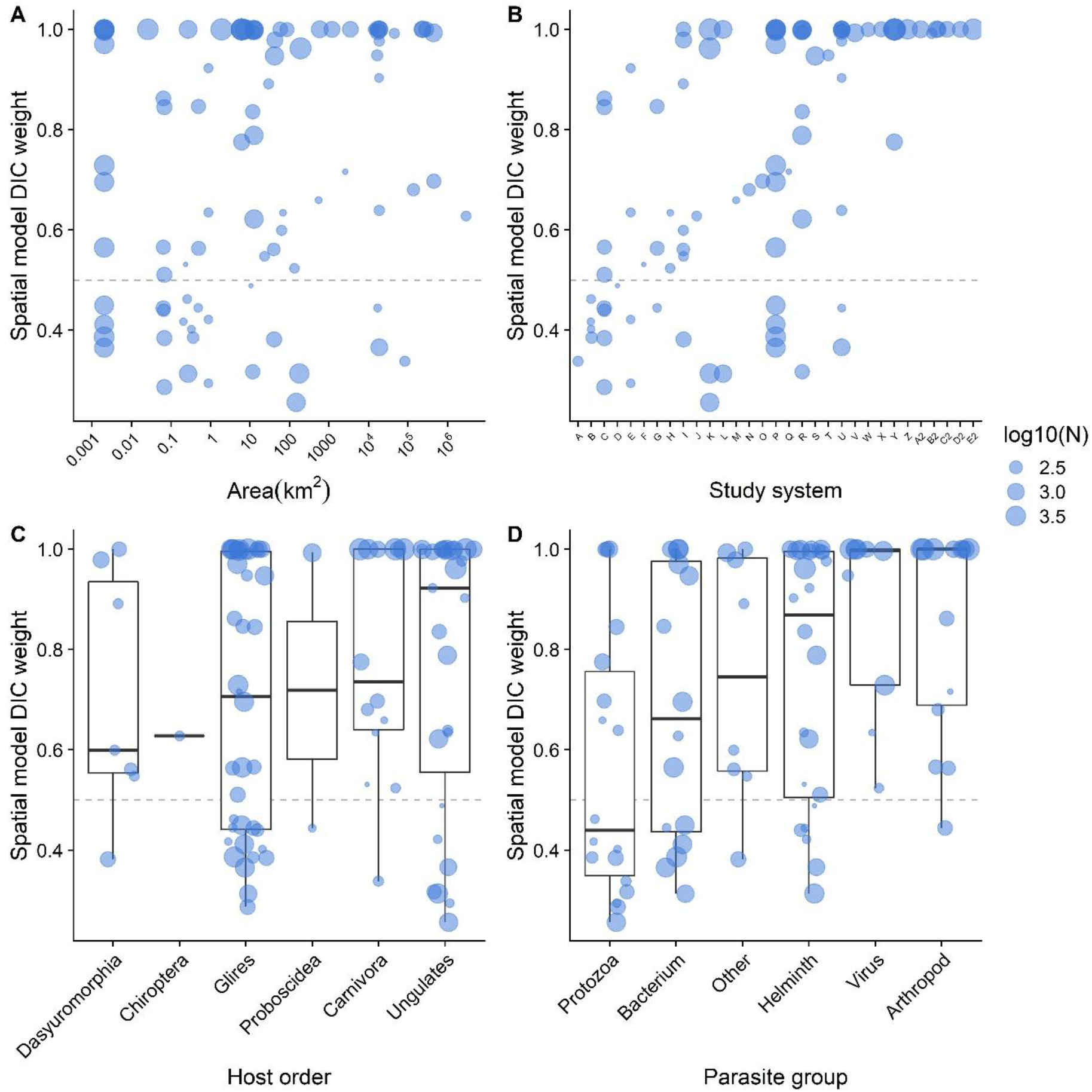
The spatial autocorrelation term (SPDE) improved models across host-parasite systems and sampling regimes. The Y axis displays the degree of confidence that the spatial autocorrelation term improved model fit (Deviance Information Criterion weight), where models at the top of the figure fitted better than those at the bottom. The dashed line at DIC weight=0.5 denotes the point at which spatial and non-spatial models were equally supported. A: larger study areas more often revealed spatial patterns. B: most of our 31 study systems exhibited at least one spatially structured host-parasite combination. Study systems have been assigned arbitrary letters to anonymise them, and are arranged in order of increasing DIC weight. C: multiple mammalian host taxa exhibited spatial effects. D: multiple parasite taxa exhibited spatial effects. The points in panels C and D are sized according to the number of samples in the replicate. None of the terms displayed here had significant effects in our meta-analysis.

Our INLA models applied across datasets consistently revealed strong spatial patterns of disease (Figure 2–3). The mean DIC change across all study systems was −14.5 (median −3.3), and the spatial model fit better than the base model for 65/89 models (73%; DIC weight>0.5). Using a conventional change of 2ΔDIC as a cutoff for improved model fit, 54% of models across 21 study systems displayed detectable spatial patterns (Figure 2). Cochran’s *Q* revealed very low heterogeneity between systems in terms of their DIC weight (*Q(df=86)* = 46.89, P=0.9998), but extreme heterogeneity in terms of the range of autocorrelation (*Q(df=86)* = 3823, P<.0001).

**Figure 3:**
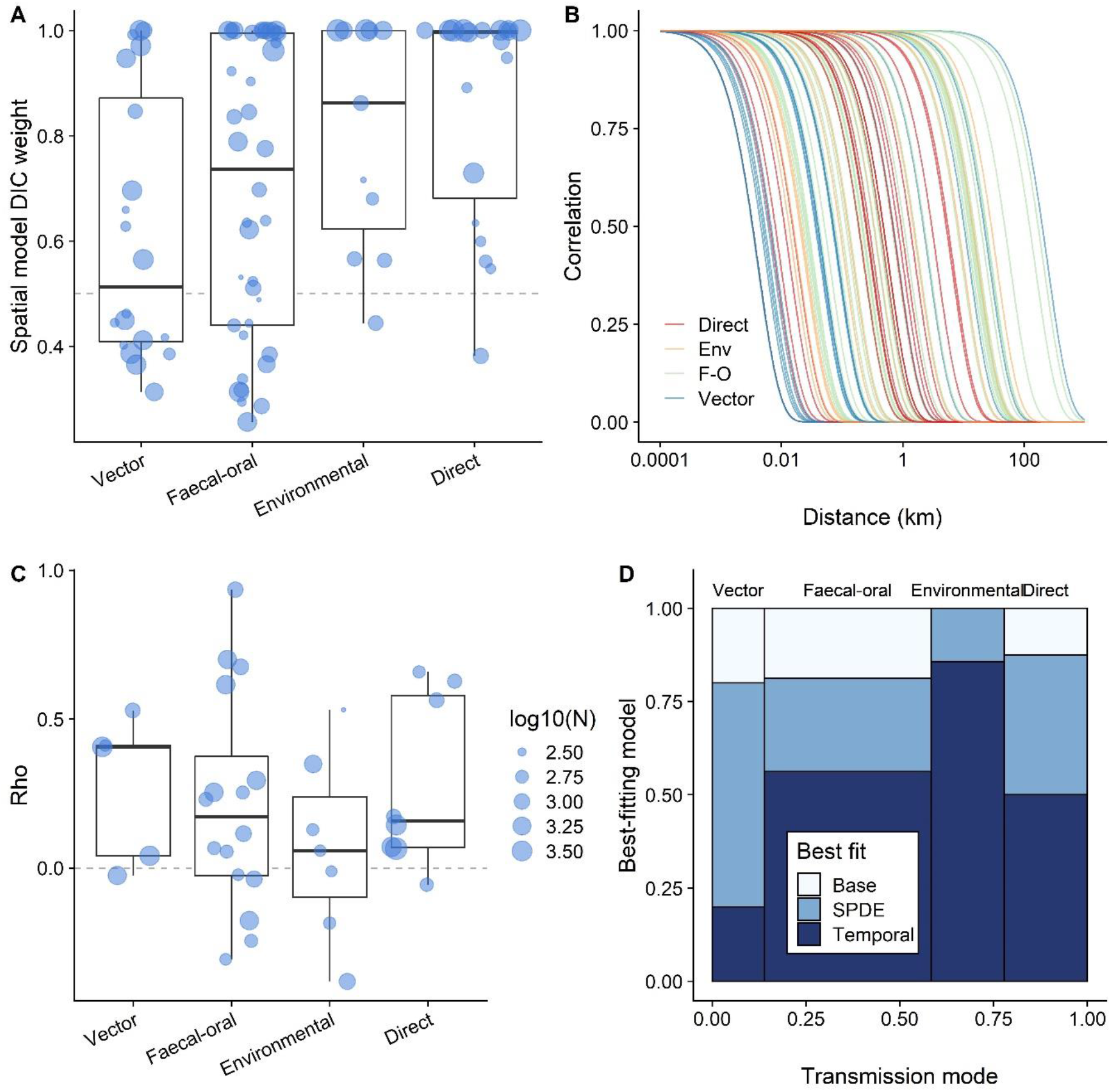
Parasites of diverse transmission modes exhibit spatial autocorrelation effects. We display A) spatial model DIC (deviance information criterion) weight, with points representing the outcome of each replicate INLA (integrated nested laplace approximation) model. Boxplots represent range, interquartile range, and median for parasites of each transmission mode. The dashed line at DIC weight=0.5 denotes the point at which spatial and non-spatial models were equally supported; points above the line display host-parasite systems for which the spatial model was better supported than the non-spatial model. B) INLA autocorrelation ranges; each line represents the autocorrelation decay of a different replicate INLA model. The colours correspond to different transmission modes, demonstrating substantial mixing of the range estimates for parasites of different transmission modes. C) Temporal autocorrelation (Rho) component demonstrating inter-annual correlations between spatial fields, for the subset of model replicates that had multiple sampling years. Points represent a different replicate INLA model; black dots represent means, and error bars represent standard errors. The dashed line at Rho=0 represents the point of no correlation; points above the line had a positive correlation, while points below the line had a negative correlation. D) Mosaic plot displaying the proportions of best-fitting models according to DIC changes, across our spatiotemporal replicates.

Although half of the systems were spatially structured, our meta-analyses revealed that few host-, parasite-, or sampling factors were predictive of spatial effects (see Supplementary Table 2). The best-fitting model for DIC weight included only the study duration (years), revealing that long-term studies were slightly more likely to uncover spatial effects (ΔAIC=3.38; for all other variables ΔAIC<1.56). The INLA range parameter increased with study area (ΔAIC=74.44) but was not affected by any other variables (ΔAIC<0.09). No variation was accounted for by host or parasite taxon, host size, or host ranging behaviour. Most notably, there was no significant variation in spatial range or DIC changes across parasite transmission modes (Figure 3A-B).

Spatiotemporal models examining a subset of multi-year studies consistently improved model fit over static equivalents. The best-fitting model for many examined replicates was a spatiotemporal model, but the findings did not differ notably across transmission modes (Figure 3D). Rho (temporal autocorrelation of the spatial field) estimates for these models were moderate, and did not vary notably across transmission modes (Figure 3C). Most (36/44, 82%) had 95% credibility intervals that overlapped with zero, and 8 (18%) were significantly positive.

## Discussion

We uncovered strong, pervasive spatial heterogeneity manifesting within an expansive diversity of mammal-parasite systems. Contrary to expectations, spatial heterogeneity was equally common and short-ranged for all transmission modes, despite our prediction that parasites with longer environmental stages would be more likely to exhibit spatial patterns. There are therefore three main takeaways from our findings: first, many study systems are spatially structured, likely by a combination of drivers, whether or not the study in question aims to quantify spatial variation or environmental drivers. Second, these drivers are relatively rarely investigated, but many systems currently have the spatial power and ability to investigate them if they wish, irrespective of the host-parasite system involved. Third, we were unable to develop a predictive framework for spatial dependence using the data available, but given more data across a wider range of host-parasite systems, such a framework may be possible to develop in the future. We therefore recommend that wild animal studies in disease ecology more regularly collect and share data on spatial behaviours and sampling locations where possible, regardless of host, parasite, or sampling regime.

Our methodology differed from that used in many other studies by investigating generalised spatial dependence rather than by quantifying specific environmental drivers that might drive this dependence. The only similar study that we know of (Gilbertson et al., 2016) used 48 parasite-locality replicates of cougar (*Puma concolor*) and bobcat (*Lynx rufus*) populations and found little evidence of spatial autocorrelation in parasite infection. In contrast to their approach, we used a wide set of different hosts, and our replicates all had between 100 and 10,000 samples (Supplementary Table 1), whereas only a few of their replicates had >100 samples, and none had >200 (Gilbertson et al., 2016). Additionally, they used Mantel tests, which do not account for fixed covariates, while the INLA analyses we employed are more suited to controlling for this variation. As such, we interpret our contrasting findings to represent a difference in the power of our analyses, and the absence of large carnivores from our dataset. Owing to its generality, similar methodology could be used in a range of ecological contexts as a useful hypothesis-generating exercise: after uncovering strong spatial structuring, researchers could follow up on this finding by investigating possible biotic or abiotic drivers. We hope that more disease ecology studies in wild animals will make use of similar methodology to bolster our understanding of disease dynamics in wild settings.

Surprisingly, neither larger study systems nor those that had previously been used to study spatial hypotheses were more likely to exhibit detectable spatial patterns. Some very small spatial replicates exhibited strong spatial effects, and the smallest area demonstrating a strong spatial trend was 0.002km^2^ (Figure 2). Similarly, some very large, well-sampled areas showed no detectable spatial patterns: anti-*Toxoplasma gondii* antibodies in almost 200 Pennsylvania black bears (*Ursus americanus*) did not (Dubey et al., 2016), while prevalence of *T. gondii* exhibited very strong spatial patterns in otters (*Lutra lutra*) across the United Kingdom (Smallbone et al., 2017), and in house mice (*Mus musculus*) within the Senegalese city of Dakar (Galal et al., 2019). However, larger study extents unsurprisingly exhibited more long-range spatial autocorrelation effects. These areas inevitably contain within them a multitude of smaller spatial effects and gradients, so that the findings of a specific study will depend critically on the spatial sampling scale it employs (Cohen et al., 2016; Luis et al., 2018; Morand et al., 2019; Pullan et al., 2012). Notably, the studies that did attempt to quantify spatial variation tended to have substantially larger spatial extent than those that did not (Supplementary Figure 2); this may represent a perception bias, where researchers operating in larger study areas tend to anticipate spatial variation as being more important to account for – or, *vice versa*, researchers asking spatial questions tend to sample across a wider range to incorporate as much testable variation as possible (Becker, Nachtmann, et al., 2019). The finding that larger study systems do not tend to more commonly exhibit detectable spatial patterns in disease demonstrates that this perception bias is perhaps unwarranted, and researchers at all scales should be able to incorporate spatial components and hypotheses about infection processes.

Despite the ubiquity of spatial effects, we discovered a very low frequency of spatial data collection and sharing: only 3 publicly available datasets included spatial data, and 22/92 responders said they had not collected any spatial data. The responses that we received implied that alongside concerns about privacy and the understandable desire to control the data associated with one’s study system, the main reason for not sharing spatial data was that the data were not collected in the first place. Location data may evade collection in some contexts where GPS signals are hard to receive, precluding spatial data collection and investigation of spatial questions. GPS instruments that function in remote environments can be expensive, and for studies that do not explicitly aim to identify spatial patterns this may seem an unnecessary expenditure. However, smartphones that can receive GPS data are now widely available and can be used in all but the most remote locations. As many researchers carry the means to collect spatial data in their pocket on a daily basis, it might take little alteration to collection protocols to include location data in many cases. Future studies should capitalise on the increasing availability of spatial telemetry and biologging technology, and associated analytical capacity (Kays, Crofoot, Jetz, & Wikelski, 2015; Long, Nelson, Webb, & Gee, 2014; Williams et al., 2020) to more frequently record, analyse, and share spatial data in disease ecology (Albery et al., 2019; Kirby et al., 2017). This practice will facilitate easier testing of the hypotheses that we outline above, as well as informing sampling regimes and mechanistic models of disease dynamics, and allowing *a priori* prediction of host-parasite systems’ spatial properties.

Privacy is an issue of considerable ethical concern in epidemiology (Kirby et al., 2017), and we contend that this concern may be contributing to a lack of open data sharing in wildlife disease ecology. Sharing spatial data risks connecting individuals with their disease status, which is particularly unwelcome in the case of stigmatised diseases such as HIV/AIDS; indeed, although we did not examine human diseases, several of the researchers we contacted opted not to share data because they were concerned that their results could be traced to specific households or individuals. Researchers could overcome this issue by jittering points, or by masking the actual GPS locations, replacing them with relative locations which are the same distance away (Kirby et al., 2017). Unfortunately, the first option will reduce precision and the latter may preclude investigation of specific geographic hypotheses or environmental drivers, but this is a small price to pay in the cases where data are potentially sensitive.

We foresee a range of potential uses for curated datasets like ours. Although we quantified some ecological and sampling-level drivers here, the dataset was still relatively small, and subject to covarying factors: for example, most analyses of nematode infection were conducted on even-toed ungulates, so that it was difficult to disentangle their implications for spatial variation. Future data collection and kind contributions from researchers may allow us to bolster this dataset to include a greater number of replicates, increasing the power and diversity of our analyses, bringing predictive models of spatial variation within our grasp. Further analysis on this dataset could investigate a number of general drivers such as population density or environmental heterogeneity, informing how they drive spatial patterns of infection within and across systems. Similar methodology could be applied to other animal groups such as birds and reptiles, whose nest and burrow locations offer ideal spatial context (e.g. Wood et al., 2007), or to marine mammals like dolphins that are regularly subject to behavioural censuses and disease surveillance (e.g. Leu, Sah, Krzyszczyk, Jacoby, & Mann, 2020). This approach could also be applied to intensively and widely spatially distributed sampling locations, either for smaller animals such as insects (Wallace et al., 2021) or for immobile organisms like plants (Halliday et al., 2020). Finally, immunity is often quantified alongside parasite burden and prevalence, and it would be interesting to see whether spatial variation in immunity manifests on the same scale, and whether it predicts disease risk (Becker et al., 2020). Given these diverse and widespread opportunities, popularising the breadth and frequency of open spatial data sharing is likely to open the door to a wide range of interesting studies and, ultimately, to the development of predictive *a priori* frameworks for spatial processes in disease ecology.

## Acknowledgements

We would like to extend our profound gratitude to the researchers who generously provided us with their data, and to all those who withstood our numerous emails regardless. GFA’s postdoctoral fellowship is funded by NSF grant number 1414296.

## Author contributions

GFA and SB conceived of the study, GFA and ARS collected the datasets, GFA conducted the INLA analyses, GFA and DJB conducted the meta-analysis, and GFA wrote the manuscript, with comments and edits offered by all authors.

## Data Availability

All code and some datasets will be made available upon publication on Github, and archived on Zenodo.org.

## Supplement

**Supplementary Table 1:** Summary table depicting the study systems used in the meta-analysis, including names, locations, host species, and sampling traits. The column “Tested spatial” denotes whether or not one of the study’s stated aims was to investigate spatial or geographic variation, e.g. in environmental drivers. The column “N” refers to the number of spatial replicates and INLA models associated with the study system. The “contributors” column refers to the individuals who generously provided data.

| Study system | Country | Location | Species | Host order | N | Parasites | Samples | Sampling method | Tested spatial? | Contributors | Example Reference |
| --- | --- | --- | --- | --- | --- | --- | --- | --- | --- | --- | --- |
| AlpineIbex | Italy | Gran Paradiso National Park | *Capra ibex* | Ungulates | 4 | Helminths; *Nematodirus*; *Marshallagia*; Coccidia | 145 | Census | N | Alice Brambilla | (Brambilla *et al.* 2015) |
| BerlinFoxes | Germany | Berlin | *Vulpes vulpes* | Carnivora | 1 | Canine distemper virus | 762 | Opportunistic | Y | Downloaded | (Gras *et al.* 2018) |
| BrazilianBats | Brazil |  | 5+ | Chiroptera | 1 | *Bartonella* | 162 | Noninvasive | Y | Keyla deSousa; Marcos Andre | Unpublished |
| BritishOtters | UK |  | *Lutra lutra* | Carnivora | 1 | *Toxoplasma gondii* | 583 | Opportunistic | Y | Derived | (Smallbone *et al.* 2017) |
| ConnecticutMice | USA | Connecticut | *Peromyscus leucopus* | Glires | 1 | Ticks | 105 | Trapping | N | Danielle Tufts; Maria Diuk-Wasser | (Tufts & Diuk-Wasser 2018) |
| ChileTriatomines | Chile |  | 5+ | Glires | 4 | Trypanosomes | 230 | Trapping | Y | Antonella Bacigalupo | (Ihle-Soto *et al.* 2019) |
| DakarRodents | Senegal | Dakar | *Mus musculus* | Glires | 1 | *Toxoplasma gondii* | 424 | Trapping | N | Lokman Galal | (Galal *et al.* 2019) |
| ElDoRodents | USA | El Dorado | 5+ | Glires | 2 | Fleas; *Bartonella* | 1612 | Trapping | Y | Michael Kosoy | (Kosoy *et al.* 2017) |
| FalkirkMice | Scotland | Callendar Park | *Apodemus sylvaticus* | Glires | 8 | *Heligmosomoides polygyrus*; *Eimeria*; *Capillaria*; Ticks; Mites; Fleas | 596 | Trapping | N | Amy Sweeny; Amy Pedersen | (Sweeny *et al.* 2019) |
| FoleyRodents | United States | California | 5+ | Glires | 3 | *Anaplasma*; *Borrelia*; Ectoparasites | 948 | Trapping | N | Janet Foley | (Foley *et al.* 2016) |
| GermanRaccoons | Germany |  | *Procyon lotor* | Carnivora | 1 | *Toxoplasma gondii* | 458 | Opportunistic | Y | Mike Heddergott | (Heddergott *et al.* 2017) |
| GreekHares | Greece |  | *Lepus europaeus* | Glires | 1 | European brown hare syndrome virus | 209 | Hunted | Y | Derived | (Sokos *et al.* 2018) |
| GrenadaMongoose | Grenada |  | *Herpestes javanicus* | Carnivora | 2 | Rabies; *Salmonella* | 157 | Trapping | Y | Ulrike Ziegler;<br>Bruno Chomel;<br>David Jaffe | (Jaffe *et al.* 2018) |
| HendyRodents | USA | California | *Neotoma fuscipes*; *Peromyscus maniculatus*; *Tamias ochrogenys* | Glires | 3 | Ticks; Aphag; Borrelia | 451 | Trapping | N | Janet Foley | Unpublished |
| HighlandDeer | Scotland | Highlands | *Cervus elaphus* | Ungulates | 1 | *Fasciola hepatica* | 103 | Hunted | Y | Andrew French | (French *et al.* 2019) |
| KielderVoles | England | Kielder Woods, Liverpool | *Microtus agrestis* | Glires | 12 | Cowpoxvirus; *Anaplasma*; *Babesia* | 3020 | Trapping | N | Mike Begon;<br>Sandra Telfer | (Davis *et al.* 2015) |
| MultimammateMice | Tanzania | Morogoro | *Mastomys natalensis* | Glires | 1 | MORV | 5547 | Trapping | Y | Lucinda Kirkpatrick | (Berkvens *et al.* 2019) |
| MyanmarElephants | Myanmar |  | *Elephas maximus* | Proboscidea | 1 | Parasite-associated mortality | 1626 | Opportunistic | N | Carly Lynsdale;<br>Virpi Lummaa | (Lynsdale *et al.* 2017) |
| PantanalMammals | Brazil | Pantanal Wetlands | *Cerdocyon thous* | Carnivora | 1 | *Hepatozoon* | 115 | Trapping | N | Marcos Andre;<br>Keyla de Sousa | (de Sousa *et al.* 2017) |
| PennsylvaniaBears | USA | Pennsylvania | *Ursus americanus* | Carnivora | 1 | *Toxoplasma gondii* | 173 | Hunted | N | Jitender Dubey;<br>Justin Brown | (Dubey *et al.* 2016) |
| RomanianFoxes | Romania |  | *Vulpes vulpes* | Carnivora | 1 | Fleas | 268 | Opportunistic | Y | Janet Foley;<br>Andrei Mihalca | (Foley *et al.* 2017) |
| RumDeer | Scotland | Isle of Rum | *Cervus elaphus* | Ungulates | 8 | Strongyles; *Nematodirus*; *Capillaria*; Coccidia; *Moniezia*; *Fasciola hepatica*; *Dictyocaulus*; *Elaphostrongylus cervi* | 2068 | Census | N | Greg Albery;<br>Josephine Pemberton;<br>Daniel Nussey | (Albery *et al.* 2019) |
| SoaySheep | Scotland | Isle of St. Kilda | *Ovis aries* | Ungulates | 4 | Strongyles; Coccidia; *Nematodirus*; Keds | 7197 | Census | N | Josephine Pemberton;<br>Daniel Nussey | (Hayward *et al.* 2014) |
| SpanishIbex | Spain | Sierra Nevada | *Capra ibex* | Ungulates | 1 | Mange | 746 | Hunted | Y | Joao Carvalho;<br>José Granados | (Carvalho *et al.* 2015) |
| StrasbourgPrimate | France | Strasbourg Primatology Centre | 5+ | Carnivora | 1 | Taeniids | 103 | Noninvasive | N | Valentin Greigert | (Greigert *et al.* 2019) |
| SwedenVoles | Sweden | Revinge | *Myodes glareolus* | Glires | 1 | *Borrelia* | 1999 | Trapping | N | Lars Raberg | (Råberg *et al.* 2017) |
| TasmanianDevils | Tasmania |  | *Sarcophilus harrisii* | Dasyuro-morphia | 7 | Devil facial tumour disease | 822 | Trapping | Y | Downloaded | (Lazenby *et al.* 2018) |
| TurnerUngulates | Namibia | Etosha National Park | *Loxodonta africana*; *Oryx gazella*; *Connochaetes taurinus*; *Antidorcas marsupialis*; *Equus quagga* | Proboscidea; Ungulates; Artiodactyla; Proboscidea | 10 | Strongyles; *Eimeria melis*; *Strongyloides* sp. | 1132 | Census | Y | Wendy Turner; Wayne Getz | (Turner *et al.* 2014) |
| WoodchesterBadgers | England | Woodchester Park, Gloucestershire | *Meles meles* | Carnivora | 1 | Bovine Tuberculosis | 3319 | Trapping | N | Matthew Silk; David Hodgson; Dez Delahay | (Rozins *et al.* 2018) |
| WythamBadgers | England | Wytham Wood | *Meles meles* | Carnivora | 4 | Fleas; Lice; Ticks; *Eimeria melis* | 7220 | Trapping | Y | Chris Newman; Christina Buesching; David MacDonald | (Albery *et al.* 2020) |
| YosemiteChipmunks | USA | Yosemite National Park, California | *Tamias speciosus* | Glires | 1 | Fleas | 1126 | Trapping | Y | Tali Hammond | (Hammond *et al.* 2019) |

**Supplementary Figure 1:**
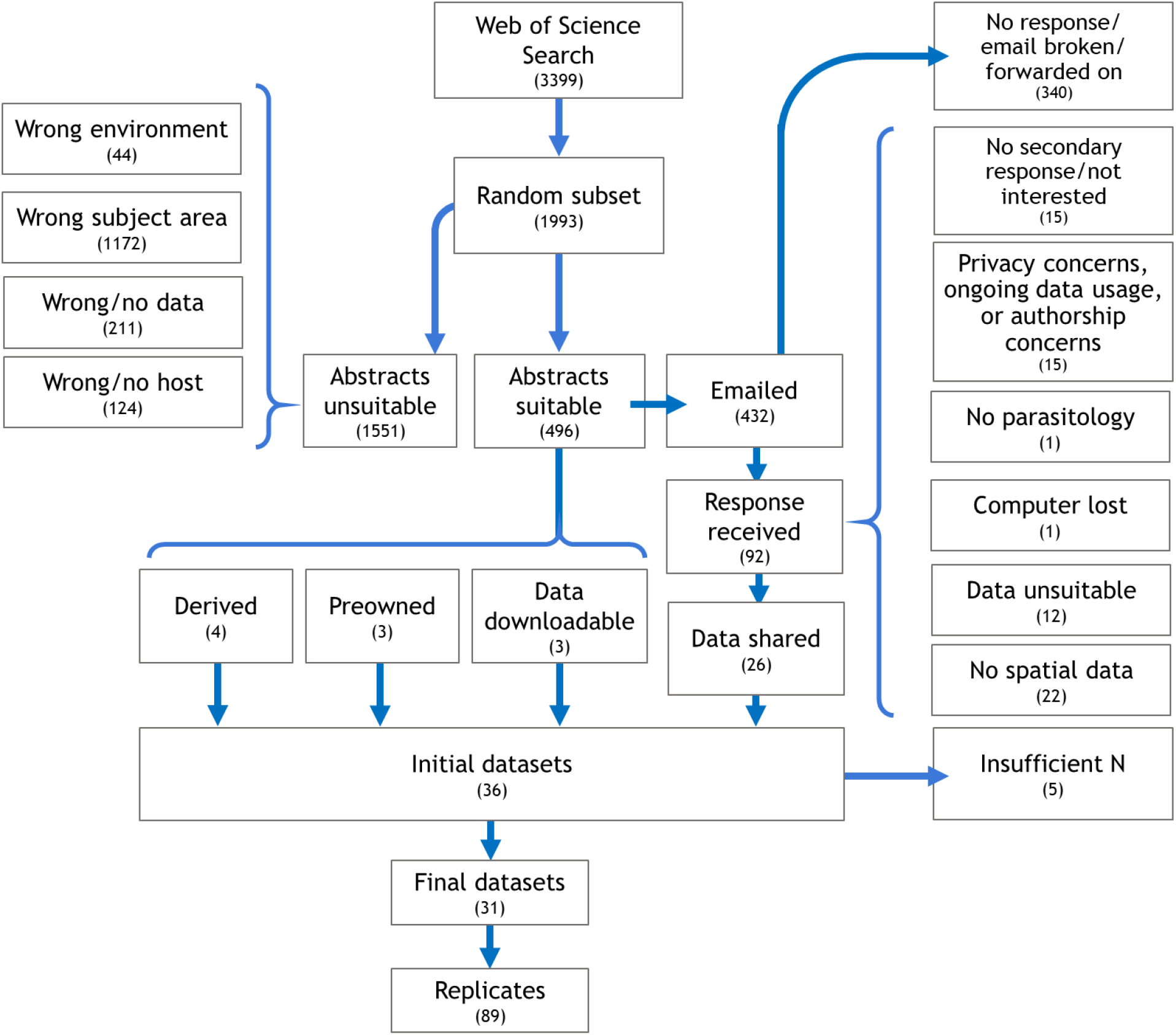
The data collection, cleaning, and analysis pipeline.

**Supplementary Figure 2:**
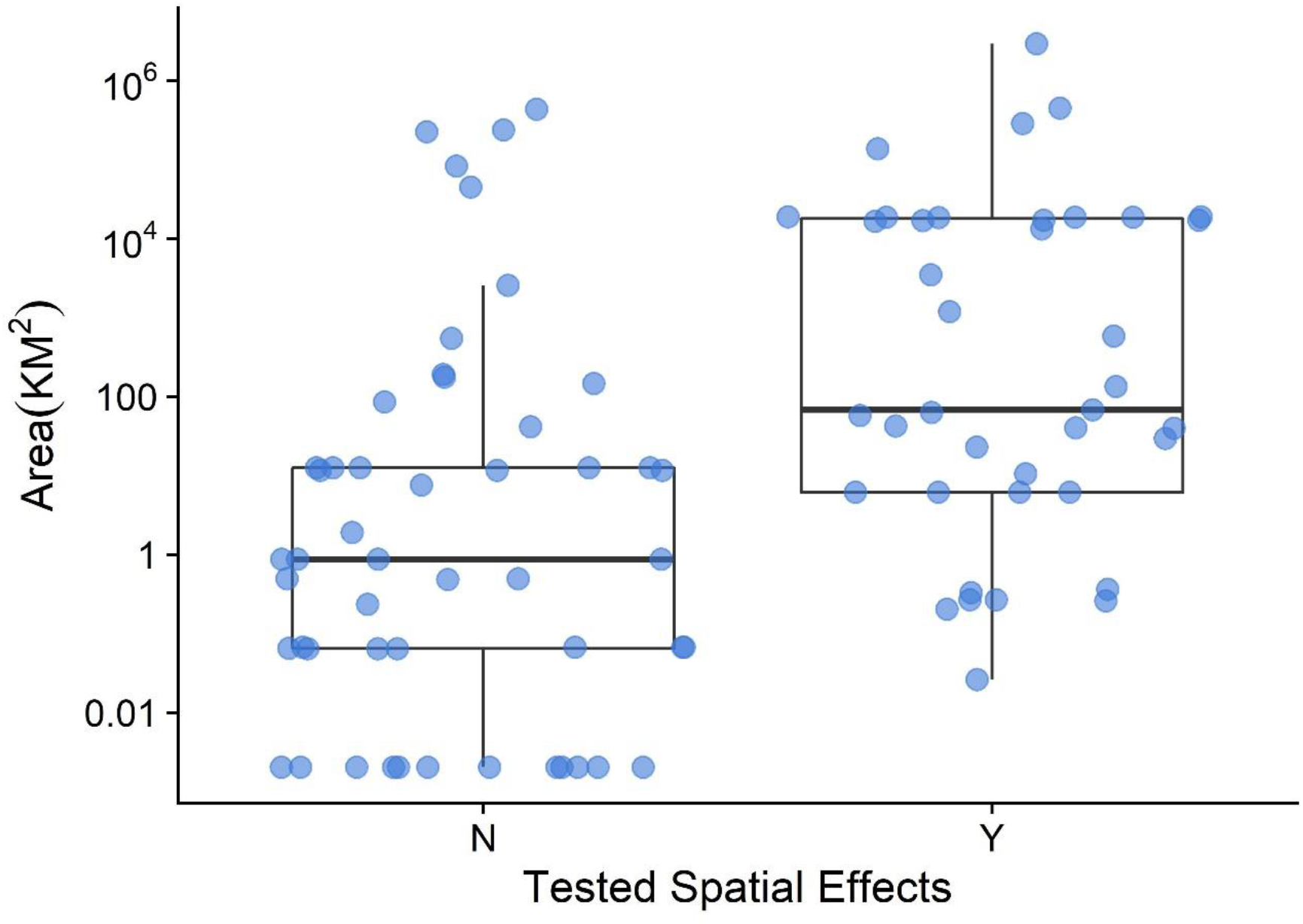
Datasets that were previously used to test spatial hypothesis were larger. The y axis is in log-km^2^. The boxplots represent the range, the interquartile range, and the median.

**Supplementary Table 2:**
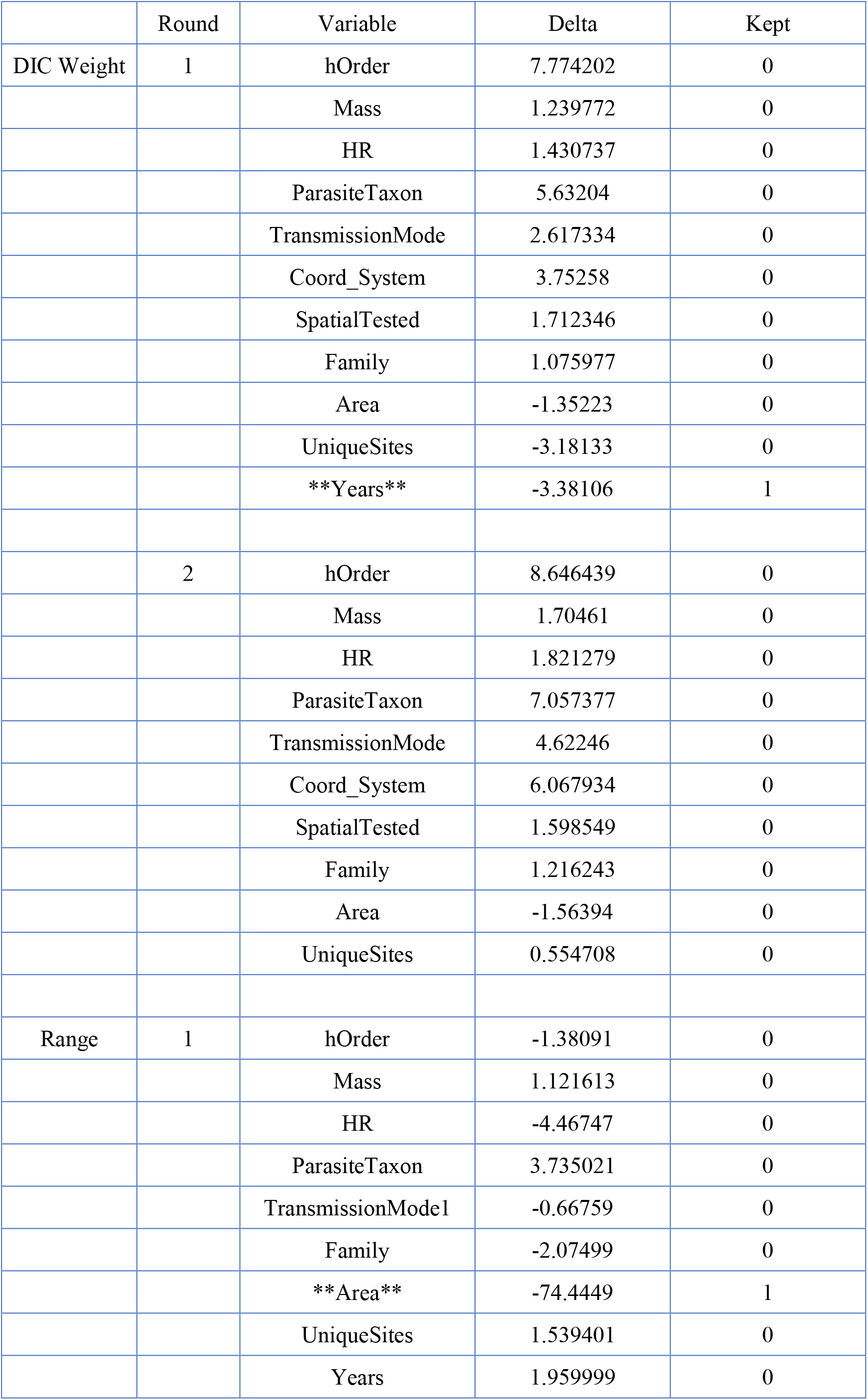
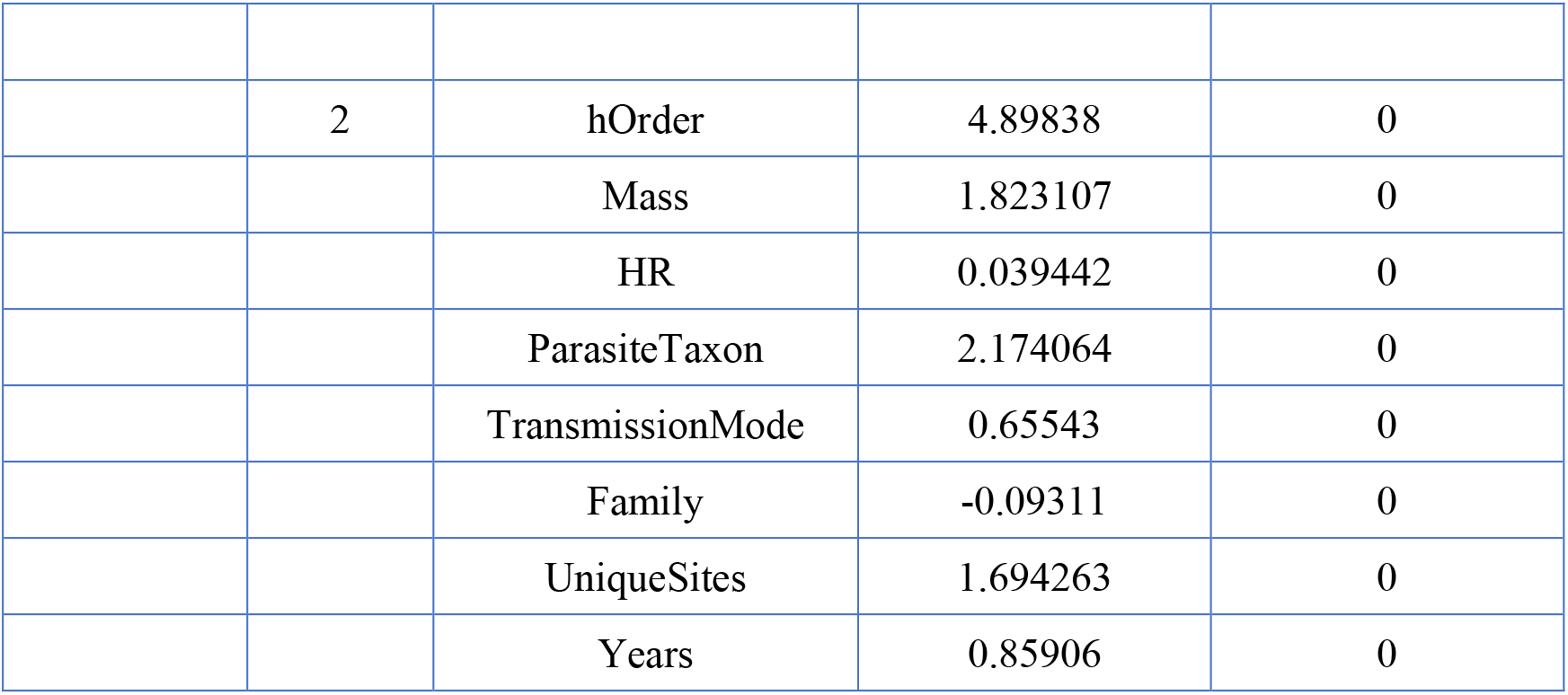
Akaike Information Criterion (AIC) changes associated with variable addition to meta-analysis models. Variables with asterisks were kept to the following round of model addition.

